# Development of the terminal airspaces in the Gray short-tailed opossum (*Monodelphis domestica*) – 3 D reconstruction by microcomputed tomography

**DOI:** 10.1101/2023.09.22.559048

**Authors:** Kirsten Ferner

**Affiliations:** Museum für Naturkunde, Leibniz-Institut für Evolutions- und Biodiversitätsforschung, Invalidenstraße 43, 10115 Berlin, Germany

**Keywords:** surface area, marsupial, morphogenesis, µCT, lung

## Abstract

Marsupials are born with structurally immature lungs when compared to eutherian mammals. The Gray short-tailed opossum (*Monodelphis domestica*) is born at the late canalicular stage of lung development. Despite the high degree of immaturity, the lung is functioning as respiratory organ, however supported by the skin for gas exchange during the first postnatal days. Consequently, the majority of lung development takes place in ventilated functioning state during the postnatal period.

X-ray computed tomography (µCT) was used to three-dimensionally reconstruct the terminal airspaces in order to reveal the timeline of lung morphogenesis. In addition, lung and air space volume as well as surface area were determined to assess the functional relevance of the structural changes in the developing lung. The development of the terminal air spaces was examined in 35 animals from embryonic day 13, during the postnatal period (neonate to 57 days) and in adults. At birth, the lung of *M. domestica* consists of few large terminal air spaces, which are poorly subdivided and open directly from short lobar bronchioles. During the first postnatal week the number of smaller terminal air spaces increases and numerous septal ridges indicate a process of subdivision, attaining the saccular stage by 7 days. The 3 D reconstructions of the terminal air spaces demonstrated massive increases in air sac number and architectural complexity during the postnatal period. Between 28 and 35 days alveolarization started. Respiratory bronchioles, alveolar ducts and a typical acinus developed. With alveolarization the volume of the air spaces and the surface area for gas exchange increased markedly. The structural transformation from large terminal sacs to the final alveolar lung in the Gray short-tailed opossum follows similar patterns as described in other marsupial and placental mammals. The processes involved in sacculation and alveolarization during lung development seem to be highly conservative within mammalian evolution.

## 1. Introduction

Marsupials have a unique reproductive strategy compared to placental mammals. The early stage of development at birth and the subsequent slow postnatal development attached to the maternal teat is a characteristic feature of marsupials that distinguishes them from other mammals (Renfree, 2006). The marsupial Gray short-tailed opossum (*Monodelphis domestica*) is born approximately 13.5 days after conception in a highly immature, nearly embryonic condition. The neonate is extremely small (∼130 mg) and exhibits a minimum anatomical development possible for a newborn mammal at birth. Most of the organ systems are immature, e.g., a functioning mesonephros, liver with simple sinusoid system, cartilaginous skeleton (Ferner et al., 2017). However, in adaptation to the reproductive strategy the neonate appears to be well developed in certain respects. An advanced olfactory system, well pronounced forelimbs and a muscular brachial plexus allow the neonate to crawl from the vagina to the mammary patch, attach itself to a maternal teat and start to suckle immediately.

Compared to eutherian neonates, marsupials are born generally with structurally immature lungs at the canalicular or saccular stage (Hill and Hill, 1955; Krause and Leeson, 1975; Gemmell and Little, 1982; Gemmell, 1986; Gemmell and Nelson, 1988; Runciman et al., 1996, 1998; Frappell and Mortola, 2000; Makanya et al., 2001, 2003, 2007; Burri et al., 2003; Szdzuy et al., 2008; Simpson et al., 2013; Modepalli et al., 2018; Ferner, 2021a). The Gray short-tailed opossum is born with lungs at the canalicular stage of lung development. Consequently, the majority of lung development occurs postnatally in air attached to the maternal teat.

While the lung in marsupials appears structurally immature, it shows qualitative characteristics of a mature gas-exchanging organ, e.g., surfactant (Ribbons et al., 1989; Miller et al., 2001; Makanya et al., 2007), a thin blood-gas barrier (Runciman et al., 1996; Szdzuy et al., 2008), neuronal-muscular reflex control of breathing (Frappell and MacFarlane, 2006). Thus, from the viewpoint of passive mechanics there might be no major constraints to inspiration (Makanya et al., 2007). However, poor muscle co-ordination and chest-wall distortion cause severe constraints to pulmonary ventilation (MacFarlane and Frappell, 2001). These neural and mechanical constraints at birth necessitate recruitment of an alternative organ system such as the skin for gas exchange (Mortola et al., 1999; Frappell and Mortola, 2000; MacFarlane and Frappell, 2001; MacFarlane et al., 2002; Frappell and MacFarlane, 2006). Cutaneous respiration is enabled by a subcutaneous capillary network with low air-blood diffusion distances, a large surface area to volume ratio, low metabolic rate and the presence of cardiac shunts in these immature newborns (Runciman et al., 1995; Frappell and Mac Farlane, 2006; Simpson et al., 2013; Ferner, 2018, 2021b). Even though supported by cutaneous respiration, most marsupials have functioning lungs at birth and rely on them as major gas exchanging organ.

The mammalian lung development was studied mainly in eutherian species, e.g., in rats, and can be categorized into five morphological stages (embryonic, pseudoglandular, canalicular, saccular and alveolar) based on characteristic morphology (Post and Copland, 2002; Tschanz, 2007; Warburton et al., 2010; Schittny, 2017).

Lung development starts in the embryonic stage (prenatal days E11–E13 in rats) with the formation of the two lung buds. At the terminal ends of the buds, a repetitive process starts where elongation of the future airways alternates with branching. The major airways and the pleura are formed. In the pseudoglandular stage (E13–E18.5 in rats) the preacinar branching pattern of airways and blood vessels is established (Kitaoka et al., 1996; Jeffery, 1998). The following canalicular stage (E18.5–E20 in rats) is characterized by branching of the terminal bronchi, terminating in small canaliculi and differentiation of type I and type II alveolar epithel cells (AECs). Towards the end of this period, the terminal or acinar tubes narrow and give rise to small saccules. Epithelial differentiation and angiogenetic activation of the capillaries lead to the first functional air-blood barriers in the lung (Frappell and MacFarlane, 2006; Morrisey and Hogan, 2010; Haberthür et al., 2021). The development of the lung proceeds with the saccular stage (E20 to postnatal day 4 in rats), which is characterized by saccule expansion, tissue proliferation, septal development and remodeling. During the alveolar stage respiratory airways and acini develop (Branchfield et al., 2016). The gas-exchange area is further enlarged by lifting off new septa from the existing gas-exchange surface and subdivision of the terminal air spaces (Burri et al., 1974; Schittny, 2017). During microvascular maturation, the double-layered capillary network of the alveolar septa is reduced to a single-layered one to increase the efficiency of the lung (Burri, 1974, 1997, 2006).

Alveolarization can be divided into two distinct phases and continues in the postnatal period (Schittny et al., 2008; Tschanz et al., 2014). During classical alveolarization (postnatal day 4– 21 in rats), new septa are formed from preexisting immature septa which contain a double-layered capillary network. During continued alveolarization (P14 to approximately P60 in rats), new septa are formed from preexisting mature single-layered capillary septa.

In eutherians, the majority of lung development occurs throughout intrauterine life. The lungs of most newborn eutherians are at the alveolar stage and the key changes that occur during early postnatal life include the increase of alveolar number and maturation of microvasculature (Burri, 2006; Szdzuy and Zeller, 2009; Schittny, 2017). Only very altricial eutherian neonates, such as mice, rats and shrews are born at the saccular stage, but reach the alveolar stage during the first postnatal days (Ten Have-Opbroek, 1981; Burri, 1997; Szdzuy et al., 2008).

In contrast to eutherian neonates, marsupials go through most of their lung development in the postnatal period. The developmental degree of the lung in newborn marsupials corresponds to the Carnegie stage 16-17 in the human fetus or E13-E14 in the fetal rat (Wang et al., 2009).

In recent years the establishment of Micro-CT techniques in combination with 3-D remodeling allowed to examine the three-dimensional structure of the lung and provided insight into the alveolarization of mouse and rat lungs (e.g., Mund et al., 2008; Schittny et al., 2008; Vasilescu et al., 2012; Haberthür et al., 2021). So far, only one study examined the developing lung of two marsupial species by phase contrast imaging methods (Simpson et al., 2013). The 3-D reconstruction of the lungs revealed that only two lung sacs were present in the newborn fat-tailed dunnart, whereas the lungs of the tammar wallaby had numerous large terminal saccules. Both species underwent marked increases in architectural complexity during the postnatal period (Simpson et al., 2013).

Studies on comparative lung development in various mammalian species let assume that mammalian lung development is highly conservative and follows similar developmental pathways in all mammalian species, including marsupials and monotremes (Szdzuy et al., 2008; Ferner et al., 2009, 2017; Mess and Ferner, 2010). The stage of lung development when mammals are born is quite variable, but not the sequence of developmental steps resulting in final lung maturation. *Monodelphis domestica* resembles both the supposed marsupial and mammalian ancestor (Szdzuy and Zeller, 2009; Deakin et al., 2013; Ferner et al., 2017). Its ancestral condition and the finding that, in contrast to eutherian mammals, most of the lung development occurs postnatally in a ventilated functioning state offers a unique opportunity for a better understanding of the development of the mammalian lung. As a first step, the development of the bronchial tree in *Monodelphis domestica* was investigated (Ferner and Mahlow, 2023). The present study was targeted at the development of the terminal air spaces of the lung in the Gray short-tailed opossum during the first weeks of life using X-ray computed tomography (µCT). In addition, we aimed to obtain functional volumes of the air spaces and surface areas of the lung using three-dimensional (3D) reconstructions of computed tomography (CT) data.

## 2. Material and methods

### 2.1. Animal collection

Gray short-tailed opossums from a laboratory colony established at the Museum für Naturkunde Berlin were controlled mated for this study. The females were checked for offspring when approaching full-term (13-14 days). Young ranging from birth to 57 days post natum (dpn) and adults were collected, weighed and euthanized by anaesthetic overdose with isoflurane under animal ethics permit approved by the Animal Experimentation Ethics Committee (registration number: T0202/18). To assess possible changes around parturition, one female was euthanized short before term by day 13 of gestation and the embryos were dissected and fixed by Karnovsky fixative (Mulisch and Welsch, 2015). A total of 35 animals between 13 days post coitum (dpc) and adult were studied using X-ray computed tomography (µCT). The numbers and specifics of the specimens are given in the supplementary table. Additional eight animals (Neonat, 5, 7,14, 21, 28, 57 dpn and adult) were investigated by scanning electron microscopy (SEM). Details for these animals are provided in Szdzuy (2006).

### 2.2. Sample preparation

Early developmental stages, ranging from neonate (defined as the first 24 h at the day of birth) to 28 dpn were decapitated, to allow for lung fixation via the trachea. The whole body of the animals was immediately immersed in Karnovsky fixative (2g paraformaldehyde, 25ml distilled water, 10 ml 25% glutaraldehyde, 15 ml 0.2 M phosphate buffer) or Bouin’s solution (picric acid, formalin, 100% acetic acid, 15: 5: 1; Mulisch and Welsch, 2015) for a longer period of time and afterwards rinsed in 70% ethanol. In late developmental stages, from 35 dpn to adults the lungs were fixed by installation via the trachea and finally dissected. Karnovsky fixative was inserted in the trachea via a cannula with polyethylene catheter tubing at a pressure head of 20 cm, until fixative was emerging from nostrils and mouth. Some animals were processed for transmission and scanning electron microscopy (TEM and SEM). From neonate to 11 dpn the upper part of the trunk was cut in two halves and in older stages (14 dpn - adult) the lungs were dissected.

The specimens for electron microscopy were fixed in 2.5% glutaraldehyde buffered in 0.2 M cacodylate (pH 7.4) for 2 hours, rinsed with 0.1 M cacodylate buffer and either postfixed in 1% osmium tetroxide and embedded in epoxy resin (Araldite) for TEM or dried in alcohol (30 - 100 %), ‘critical-point-dried’, mounted, sputter-coated with gold-palladium for SEM. The samples were viewed in a scanning electron microscopic (LEO 1450 VP, Carl Zeiss NT GmbH).

### 2.3. Preparation for µCT

For visualization of the lungs in X-ray computed tomography the specimens had to be stained in advance. Different staining protocols were tested and used at different age stages (Supplementary Table). Staining with tungstic acid (PTA) was either performed in ethanol with 1% PTA for 21 to 42 days (full body specimens of 14 to 28dpn) or in an aqueous solution (Karnovsky fixative) starting with 0.5% for 7 to 20 days and increased afterwards to 1% resulting in a staining period up to 30 days (Metscher, 2009). Separated lungs of older stages and adult specimens were stained in 1% Iodine (Gignac et al., 2016) to test staining differences and shrinking effects, which could not be detected. The differences in staining periods and staining concentration depended on the respective specimen size and preparation.

The specimens processed for electron microscopy could be scanned without further processing.

### 2.4. µCT imaging

The prepared specimens were subjected to micro-tomographic analysis at the Museum für Naturkunde Berlin (lab reference ID SCR_022585) using a Phoenix nanotom X-ray|s tube (Waygate Technologies, Baker Hughes, Wunstorf, Germany; equipment reference ID SCR_022582) at 70–110kV and 75–240μA, generating 1440–2000 projections (Average 3– 6) with 750–1000ms per scan or a YXLON FF85 (equipment reference ID SCR_020917) with transmission beam for bigger specimens at 90–110kV and 100–150μA, generating 2000 projections (Average 3) with 250-500ms. The different kV, µA and projection-settings depended on the respective machine and specimen size, which is also responsible for the range of the effective voxel size between 1.5–20.1μm. The cone beam reconstruction was performed using the datos|x 2 reconstruction software (Waygate Technologies, Baker Hughes, Wunstorf, Germany; datos|x 2.2) or Nexus reconstruction software respectively.

### 2.5. 3D reconstruction

Non-destructive µCT-imaging, in particular of entire animals, offers various possibilities for different research approaches. Surface scans give an impression of the external anatomy of the examined animals (Fig. 1A-C) or the anatomical position of 3D reconstructed organs can be assessed (Fig. 1D). The 3D volume processing was carried out with the software Volume Graphics Studio Max Version 3.5 (Volume Graphics GmbH, Heidelberg, Germany). 3D X-ray tomography data were analyzed as serial two-dimensional (2D) and reconstructed to three-dimensional (3D) images (Fig.2). Since in the 2D images organ tissues were colored in different shades of gray according to the tissue’s density, formerly air-filled regions, such as the air spaces and conducting airways of the lung appeared black. The outline of the air-filled areas (terminal airspaces and bronchial tree) of the lung were reconstructed on a slice-by-slice basis and regions of interest (ROI) were created. The volumes of the bronchial trees were estimated in a previous study (Ferner and Mahlow, 2023) and could be deducted from the total air-filled volumes, resulting finally in the volumes of the terminal air spaces. The volumes and surface areas of the ROIs of the terminal air spaces were calculated by the program Volume Graphics and indicated by mm³ for volume and by mm² for surface area with an accuracy of two digits after the decimal point. The values are presented as median and range in Table 1.

**Fig. 1.**
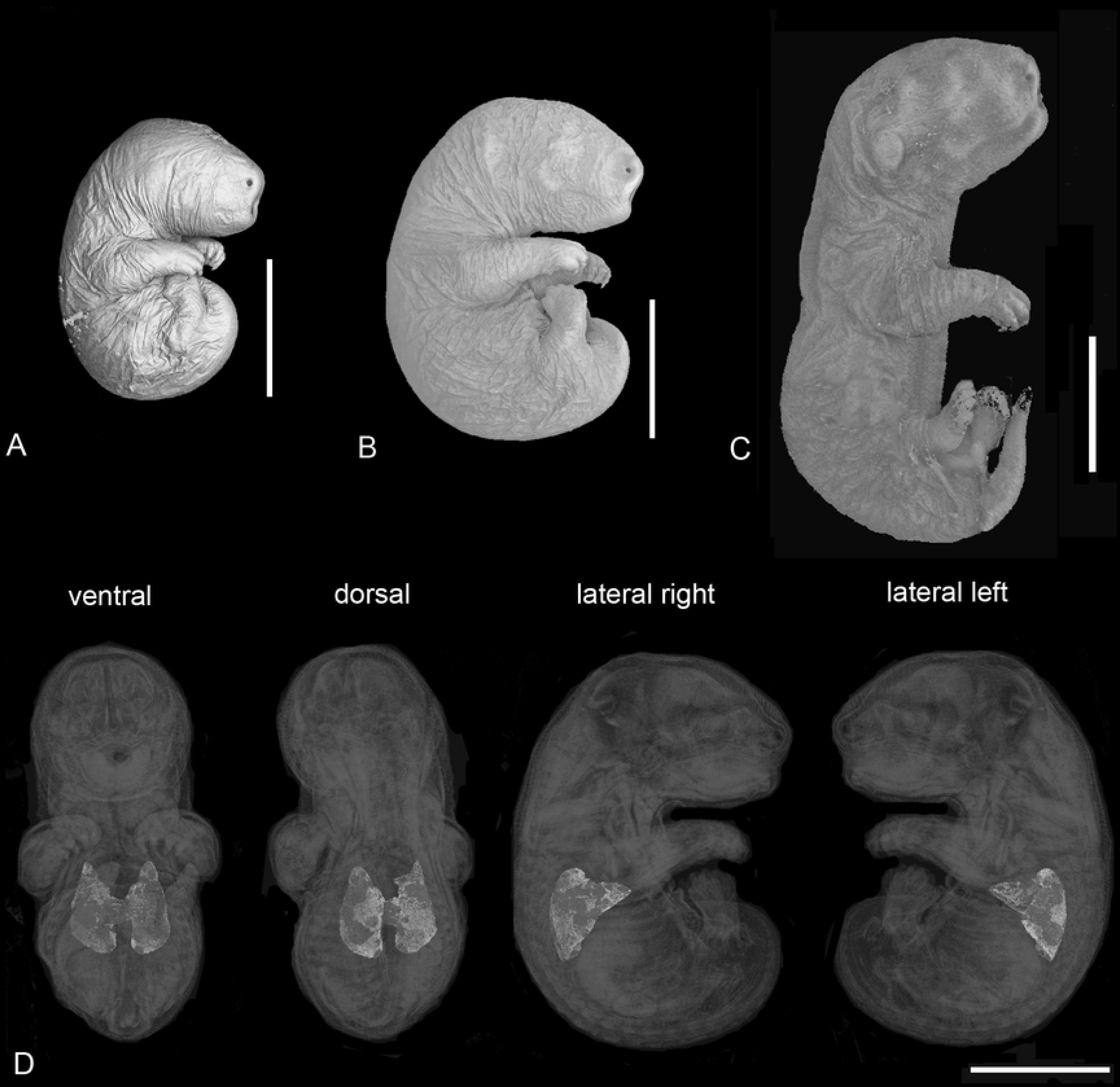
External appearance of a 4 dpn (A), 7 dpn (B) and 11 dpn (C) old Gray short-tailed opossum. Characteristic for marsupial offspring is the embryonic appearance, with stronglydeveloped forelimbs and undifferentiated paddle-like hindlimbs as well as an undifferentiated oro-nasal region with oral shield until 7 dpn. By 11 dpn the ears, oral region and hind limbs appear more differentiated. Anatomical position of the lung from ventral, dorsal and lateral views (D, from left to right). Scale bar = 5 mm.

**Fig. 2.**
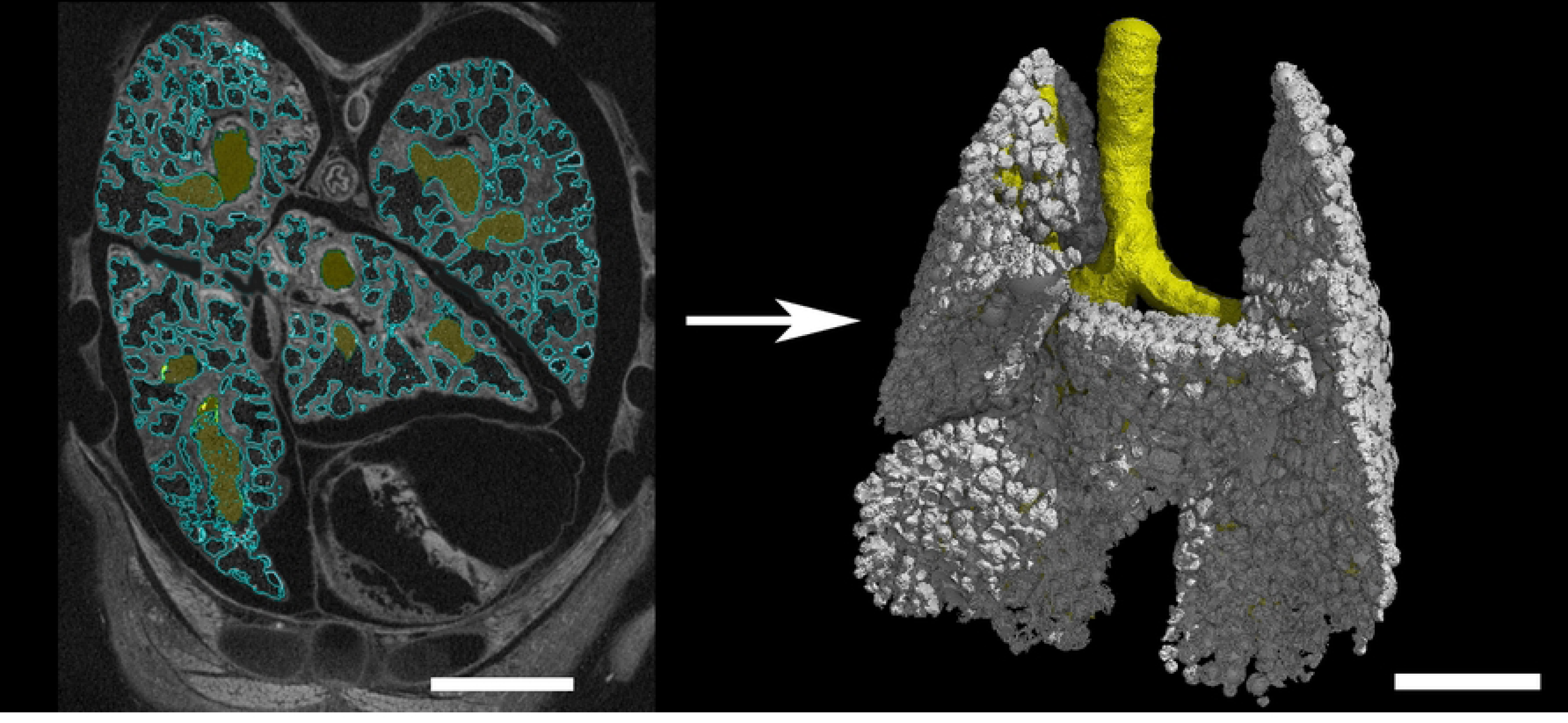
Three-dimensional reconstruction of the lung of *Monodelphis domestica* at 7 dpn (2383_2) using 3D X-ray µCT. (A) µCT image in transverse section (2D) with marked airspaces (blue) and bronchial tree (yellow). (B) 3D reconstruction of the terminal air spaces (white) and the bronchial tree (yellow). Scale bar = 0.7 mm.

**Table 1.**
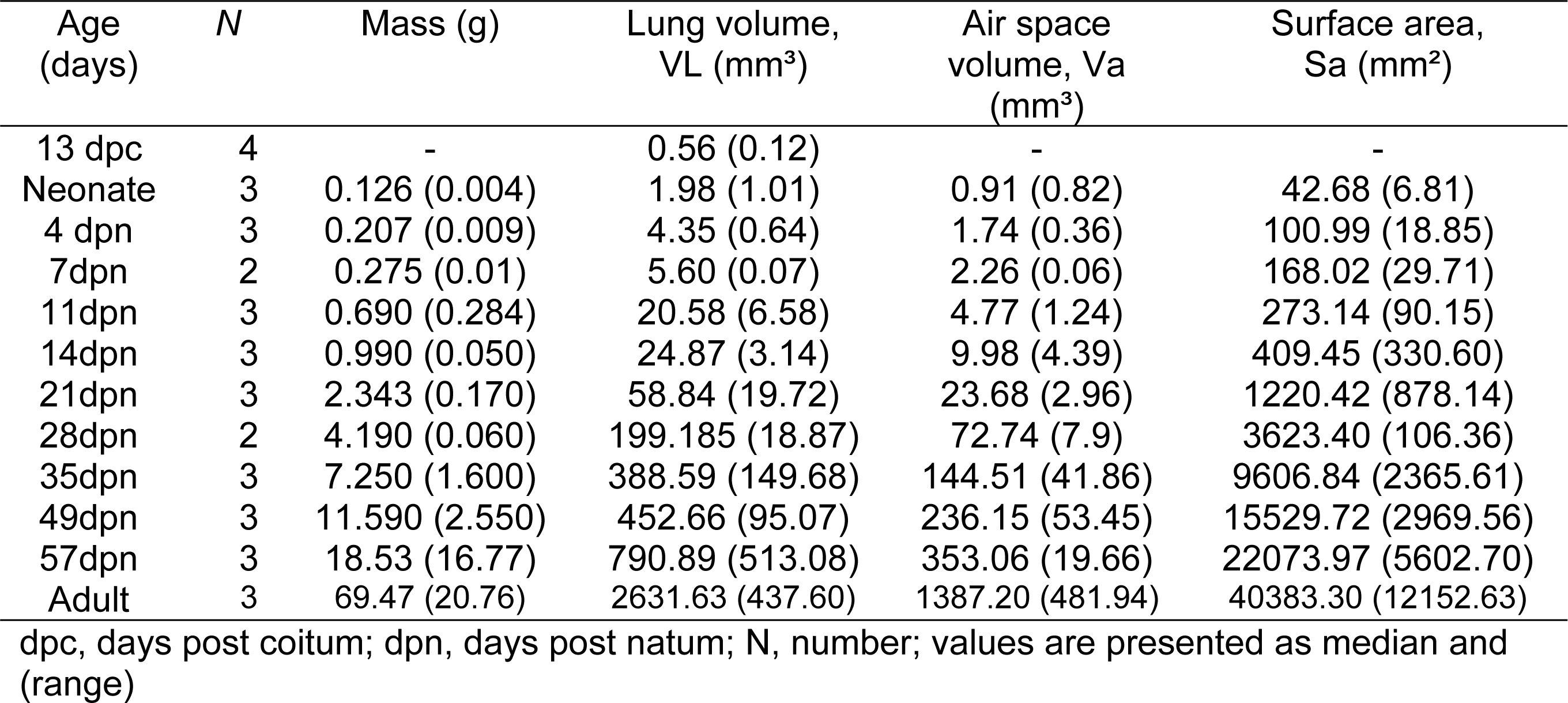
Body weights, volumes of the lung and air spaces and surface area of the Gray short-tailed opossum (*Monodelphis domestica*) during the postnatal period.

| Age<br>(days) | N | Mass (g) | Lung volume,<br>VL (mm<sup>3</sup>) | Air space<br>volume, Va<br>(mm<sup>3</sup>) | Surface area,<br>Sa (mm<sup>2</sup>) |
| --- | --- | --- | --- | --- | --- |
| 13 dpc | 4 | - | 0.56 (0.12) | - | - |
| Neonate | 3 | 0.126 (0.004) | 1.98 (1.01) | 0.91 (0.82) | 42.68 (6.81) |
| 4 dpn | 3 | 0.207 (0.009) | 4.35 (0.64) | 1.74 (0.36) | 100.99 (18.85) |
| 7dpn | 2 | 0.275 (0.01) | 5.60 (0.07) | 2.26 (0.06) | 168.02 (29.71) |
| 11dpn | 3 | 0.690 (0.284) | 20.58 (6.58) | 4.77 (1.24) | 273.14 (90.15) |
| 14dpn | 3 | 0.990 (0.050) | 24.87 (3.14) | 9.98 (4.39) | 409.45 (330.60) |
| 21dpn | 3 | 2.343 (0.170) | 58.84 (19.72) | 23.68 (2.96) | 1220.42 (878.14) |
| 28dpn | 2 | 4.190 (0.060) | 199.185 (18.87) | 72.74 (7.9) | 3623.40 (106.36) |
| 35dpn | 3 | 7.250 (1.600) | 388.59 (149.68) | 144.51 (41.86) | 9606.84 (2365.61) |
| 49dpn | 3 | 11.590 (2.550) | 452.66 (95.07) | 236.15 (53.45) | 15529.72 (2969.56) |
| 57dpn | 3 | 18.53 (16.77) | 790.89 (513.08) | 353.06 (19.66) | 22073.97 (5602.70) |
| Adult | 3 | 69.47 (20.76) | 2631.63 (437.60) | 1387.20 (481.94) | 40383.30 (12152.63) |

dpc, days post coitum; dpn, days post natum; N, number; values are presented as median and (range)

## Results

The lungs of the Gray short-tailed opossum consist of six lung lobes, a cranial, a middle, a ventral and a caudal lobe in the right lung and a middle and a caudal lobe in the left lung. Figure 3 shows the ventral, dorsal, lateral, cranial and caudal views of the lung lobes in the newborn *Monodelphis domestica*. The lungs of the near-term fetus at 13 dpc and in the neonate are at the canalicular stage of lung development and consist of large terminal air sacs, which open directly from the lobar bronchioles (Fig. 4 A, B; 5 A-C, D-F). The terminal air spaces are deflated before birth, yielding a low lung volume of 0.56 mm³. With birth the lungs become ventilated and the conducting airways and terminal airspaces are expanded by air, resulting in a lung volume of 1.98 mm³ in the neonate (Tab.1). At this time the gas exchange takes place in the distal portions of the conducting airways, which are lined with respiratory epithelium, and the large terminal air spaces, which have a lumen of ∼ 450 µm in diameter and are subdivided only a little (Fig. 5F, 6A). A thick interstitial layer (∼45 µm) separates the capillaries of one air space from the capillary network of the adjacent air space resulting in a thick septum (Fig. 6A).

**Fig. 3.**
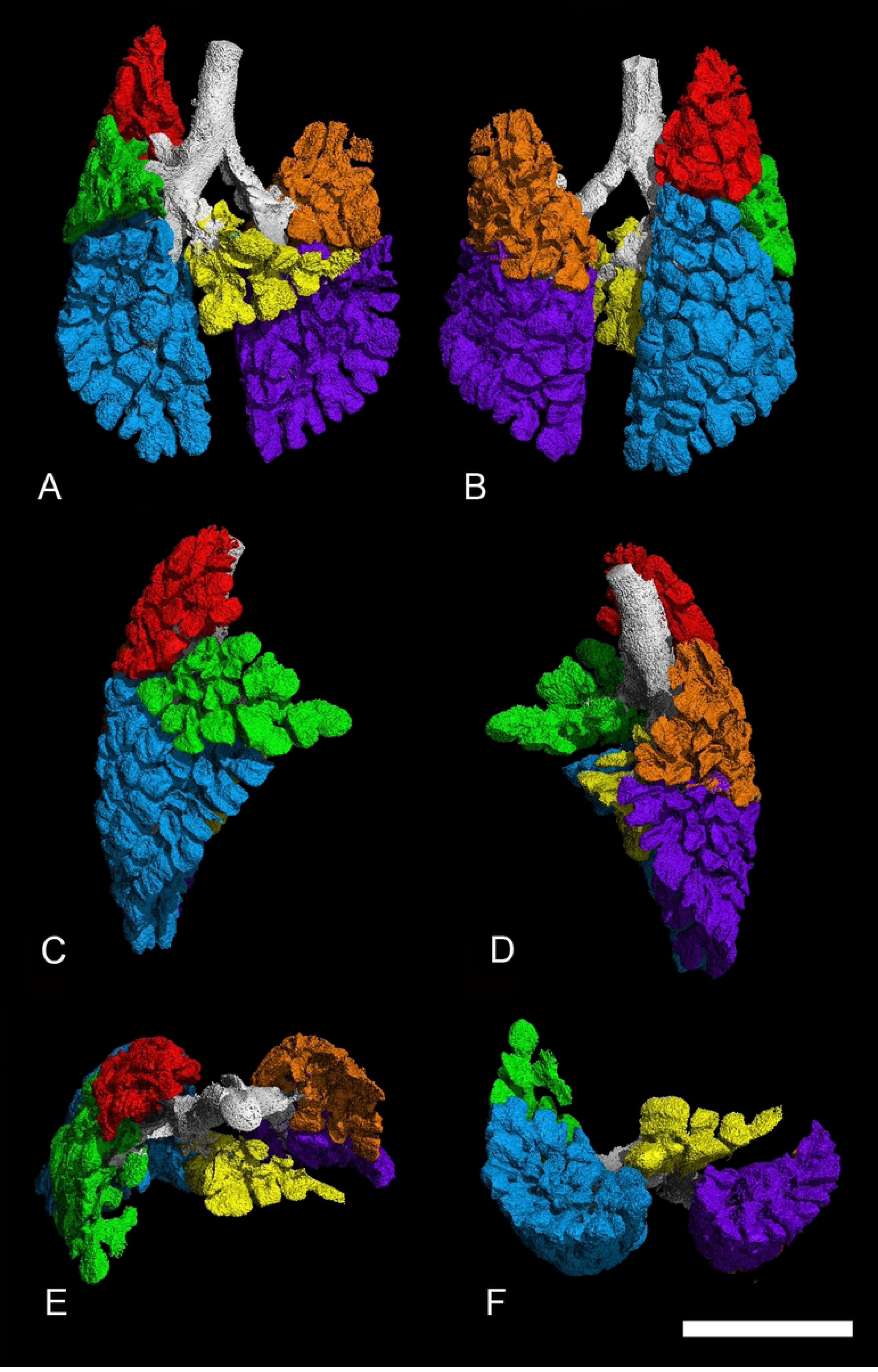
Reconstruction of the terminal airways in the newborn lung of Monodelphis domestica with differentiation of the pulmonary lobes. The lungs are shown in ventral (A), dorsal (B), lateral views from the right (C) and left (D) side and perspectives from the cranial (E) and caudal side (F). The pulmonary lobes are indicated by colors: right lung – cranial lobe (red), middle lobe (green), accessory lobe (yellow) and caudal lobe (blue); left lung – middle lobe (orange) and caudal lobe (purple). Scale bar = 1 mm

**Fig. 4.**
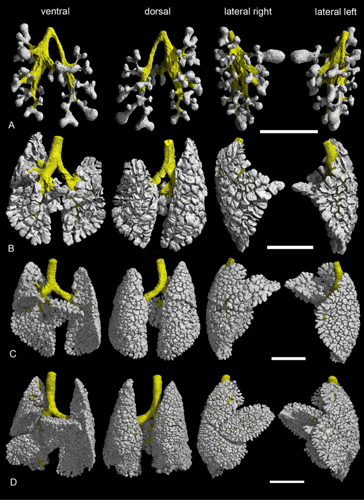
Representative 3D reconstructions of the terminal airspaces of *Monodelphis domestica* at 13 dpc (A), in the neonate (B), at 4 dpn (C) and at 7 dpn (D). The lungs are shown from different perspectives: in ventral, dorsal and lateral views from the right and left side (from left to right). The scale bar is 0.5 mm in (A) and 1 mm in (B-D).

**Fig. 5.**
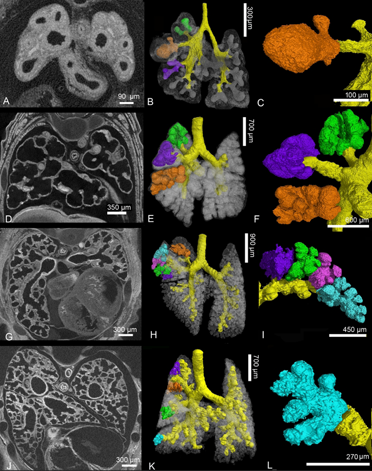
Details of the developing *Monodelphis domestica* lung at 13 dpc (A-C), neonate (D-F), 4 dpn (G-I) and 7 dpn (J-L). In embryos of 13 dpc and in neonates large terminal air spaces branch off directly from a simple bronchial tree, each forming a pulmonary lobe (F). By 4 dpn the terminal air spaces become subdivided by septal growth. Branching from new formed segmental bronchioles several terminal air spaces can be distinguished in the pulmonary lobes (I). By 7 dpn the sub segmentation of the air spaces progresses and the terminal saccules decrease in size (L). 2 D sections: A, D, G, J; position of terminal airspaces in the lung: B, E, H, K; close-up view of terminal air spaces: C, F, I, L. The Magnification is indicated by the scale bar.

**Fig. 6.**
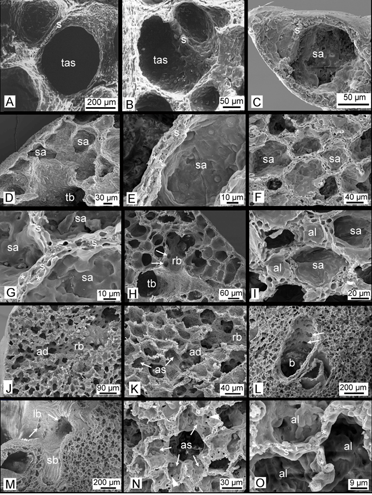
Scanning electron micrographs of the neonate (A); 4 dpn (B), 7 dpn (C), 14 dpn (D, E), 21 dpn (F, G), 28 dpn (H, I), 57 dpn (J-L) and adult (M-O) Monodelphis domestica lung. The newborn and 4 dpn lung are characterized by large terminal airspaces, which become successively smaller with progressing subseptation. By 14 dpn and 21 dpn numerous small sacculi, which are separated from each other by thin double capillary septa (capillary beds are indicated by asterisks). By 28 dpn first alveoli (indicated by arrows) can be seen and respiratory bronchioles develop. By 57 dpn and in adults a mature lung with respiratory bronchioles, alveolar ducts and alveolar sacs is present. Between alveoli single capillary septa have been formed. ‘Pores of Kohn’ (indicated by arrowheads) form interalveolar connections in walls of adjacent alveoli and connect them to each other. Magnification is indicated by the scale bar. ad, alveolar duct; al, alveolus; as, alveolar sac; b, bronchiole; lb, lobar bronchiole; rb, respiratory bronchiole; s, septum; sa, sacculus; sb, segmental bronchiole; tas, terminal air spaces; tb, terminal bronchiole. The Magnification is indicated by the scale bar.

The lung of a four days old *Monodelphis domestica* distinguishes from that of a newborn particularly through an increasing subdivision of the terminal air spaces (Fig. 4C, 5G-I, 6B). The terminal air sacs are still large with a diameter of 200-300 µm, but a number of smaller saccules and septal ridges indicate a process of subdivision of the terminal air spaces. The septa separating the terminal air sacs vary in thickness. Those in the central part of the lung are thicker (40 µm) than the more peripherally located septa (20 µm).

By 7 dpn a further subdivision of the terminal airspaces and a gradually decrease in size of the terminal air sacs can be seen (Fig. 4D, 5J-L, 6C). A continuous double capillary bed, however with a thick interstitial layer, is present in the septa and indicates the transition to the saccular stage of lung development. The terminal air sacs have a size of 150-200 µm in diameter. The new formed smaller terminal air sacs appear smooth walled, whereas crests that vary in height and thickness are numerous in the larger air sacs, resulting in an irregular shape.

Further compartmentalization occurs in the lung between 11 and 21 postnatal days (Fig. 7A-C). Substantial changes take place in the architecture of the lung (Fig. 8A-L). The terminal saccules become more numerous and decrease in size (Fig. 6D, F). They measure ∼150 µm in diameter by 14 dpn and ∼90 µm by 21 dpn. Several new saccules develop near the pleura. They are separated by septa vertically standing on the pleura (Fig. 8K). The saccules are still separated by a double capillary septum (Fig. 6E, G). However, the septa decrease in thickness and measure ∼15 µm by 21 postnatal days. In contrast to earlier stages a thick core of stromal cells with large interstitial spaces is missing (Fig. 6G). The lung volume and surface area improve through extensive increases in saccular number, as well as architectural complexity (Tab. 1).

**Fig.7.**
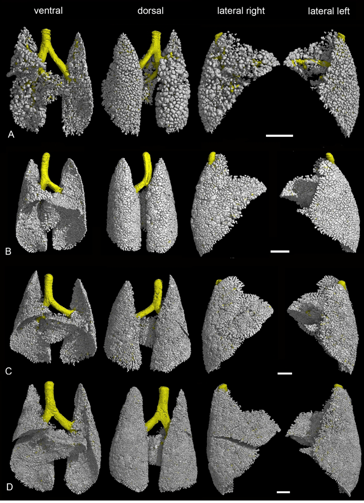
Representative 3D reconstructions of the terminal airspaces of *Monodelphis domestica* at 11 dpn (A), 14 dpn (B), 21 dpn (C) and at 28 dpn (D). The lungs are shown from different perspectives: in ventral, dorsal and lateral views from the right and left side (from left to right). The scale bar 1 mm.

**Fig. 8.**
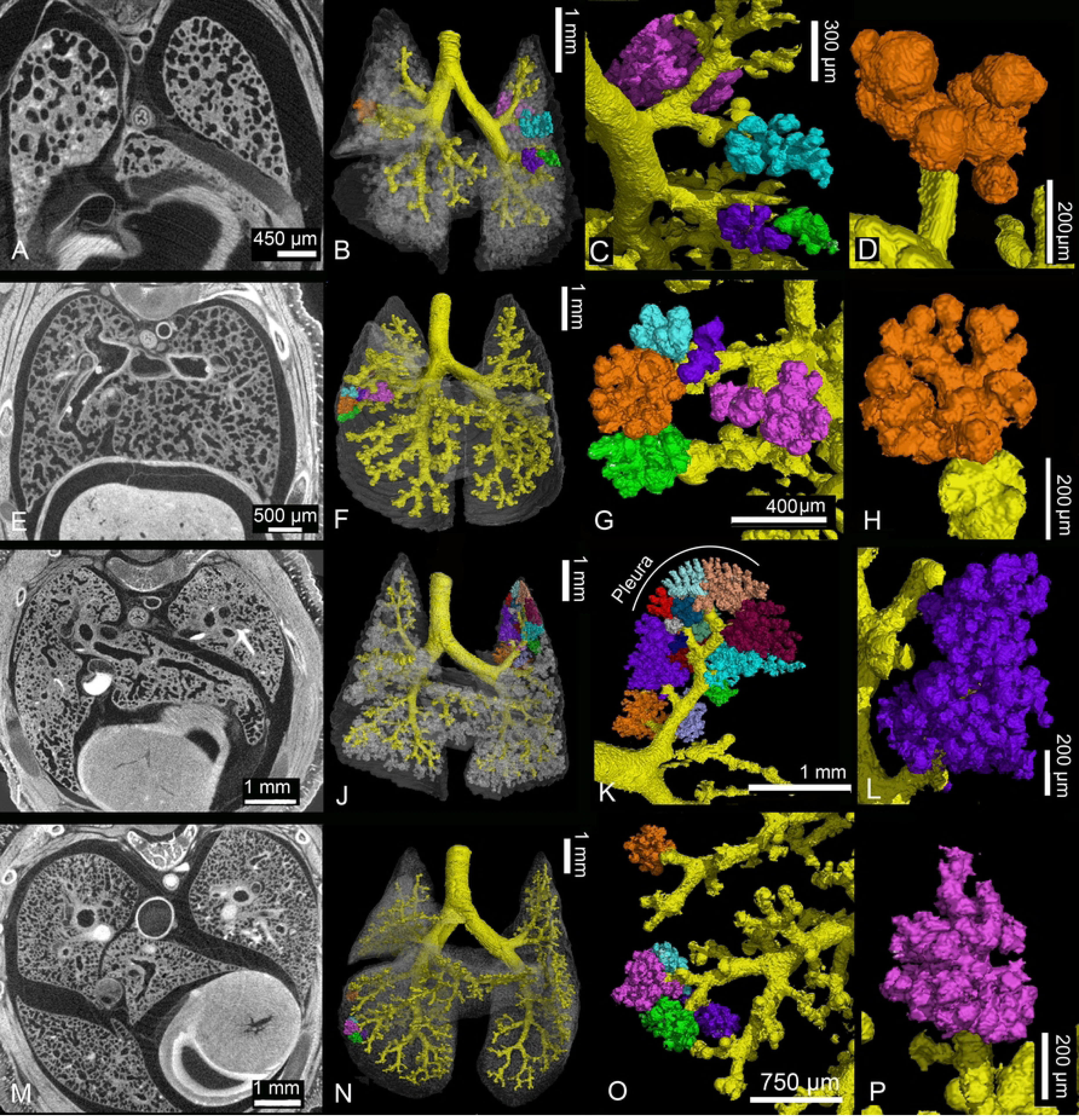
Details of the developing *Monodelphis domestica* lung at 11 dpn (A-D), 14 dpn (E-H), 21 dpn (I-L) and 28 dpn (M-P). Compartmentalization of the terminal air spaces progresses between 11 and 21 postnatal days. The terminal saccules become more numerous and decrease in size. Several new saccules develop near the pleura, separated by septa vertically standing on the pleura (8K, red, bright blue and beige). By 28 dpn alveolarization starts. The terminal air spaces consist of saccules and new formed alveoli. 2 D sections: A, E, I, M; position of terminal airspaces in the lung: B, F, J, N; terminal air spaces: C, G, K, O; close-up view of terminal air space: D, H, L, P. The Magnification is indicated by the scale bar.

With 28 days the transition from the saccular to the alveolar period of lung development begins. The terminal air spaces, consisting of saccules and new formed alveoli, are characterized by a further increase in number and decrease in size (Fig. 7D, 8M-P). Saccules still dominate the lung parenchyma by 28 dpn. Besides the larger saccules (∼60 µm), which are separated from each other by a double capillary septum, a few smaller alveoli (40-50 µm), separated by a single capillary septum, can be seen (Fig. 6I). Compared to the earlier developmental stages, the thickness of both, the double and single capillary septa, decreases furthermore. They measure ∼13 µm for double capillary septa and 6-8 µm for single capillary septa. With the formation of alveoli first respiratory bronchioles develop. They are characterized by flattened epithelium with alveoli located in their walls (Fig. 6H). The further increase in saccular number and new formation of alveoli lead to an increase in lung volume, air space volume and surface area by 28 dpn (Tab. 1).

The lung of the 35 days old *Monodelphis domestica* is characterized by a further increase in lung volume and the proceeding formation of alveoli (Tab.1, Fig. 9A, 10A-D). The lung has fully attained the alveolar stage. The bulk of alveoli lead to a further increase in air space volume and surface area (Tab. 1). Respiratory bronchioles with alveoli are found more frequently compared to 28 dpn. The distal parts of the respiratory bronchioles pass into alveolar ducts which open into alveolar sacs. Thus, typical structures of the mammalian acinus are present.

**Fig. 9.**
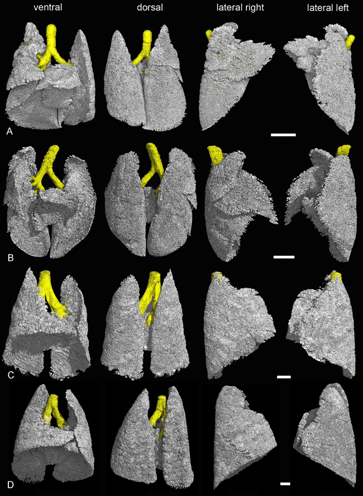
Representative 3D reconstructions of the terminal airspaces of *Monodelphis domestica* at 35 dpn (A), 49 dpn (B), 57 dpn (C) and in an adult (D). The lungs are shown from different perspectives: in ventral, dorsal and lateral views from the right and left side (from left to right). The scale bar 2 mm.

**Fig. 10.**
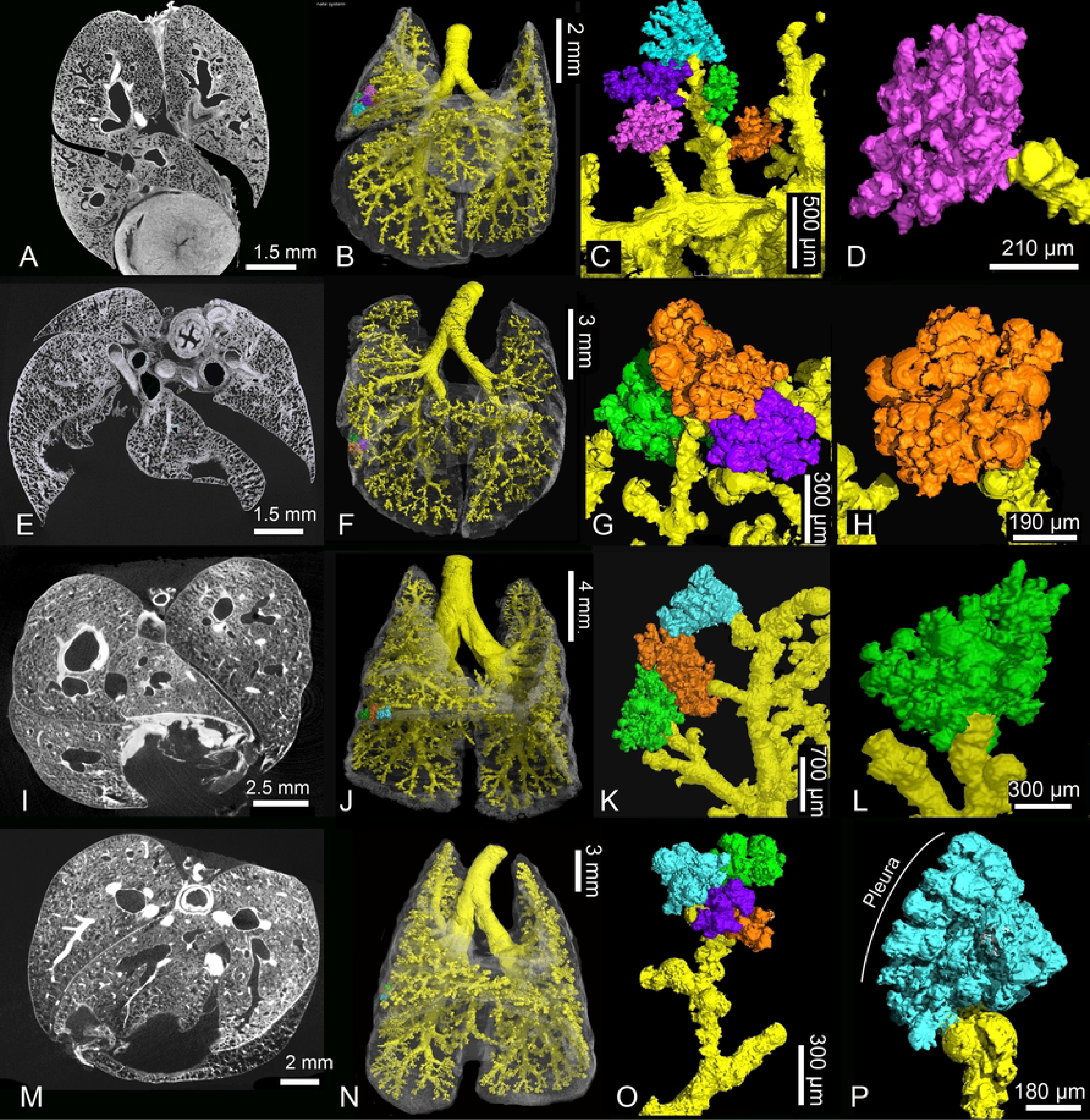
Details of the developing *Monodelphis domestica* lung at 35 dpn (A-D), 49 dpn (E-H), 57 dpn (I-L) and adult (M-P). By 35 dpn the lung has fully attained the alveolar stage. The terminal airways consist of respiratory bronchioles and alveolar ducts. The lung parenchyma at 49 dpn, 57 dpn and in adult *Monodelphis domestica* is strongly subdivided and the alveoli in their entirety provide a large surface area for gas exchange. Alveoli located at the pleural surface are separated by septa vertically standing on the pleura (P) similar as observed in sacculi close to the pleural surface by 21 dpn. 2 D sections: A, E, I, M; position of terminal airspaces in the lung: B, F, J, N; terminal air spaces: C, G, K, O; close-up view of terminal air space: D, H, L, P. The Magnification is indicated by the scale bar.

By 49 dpn (Fig. 9B, 10E-H) no distinct structural changes can be seen in the terminal airspaces of *Monodelphis domestica*. In the 57 days old lung (Fig. 9C, 10I-L) alveoli have markedly increased in number. With the progressing formation of alveoli, numerous respiratory bronchioles can be seen (Fig. 6J). The distal parts of the respiratory bronchioles pass into short alveolar ducts, which are covered with respiratory epithelium and have alveoli at their sides. The alveolar ducts open into alveolar sacs, from which alveoli radiate into the surrounding parenchyma (Fig. 6K). The alveoli measure ∼40 µm in diameter. In addition to this regular development, alveoli are present also at the walls of solely conducting airways, such as segmental bronchi and terminal bronchiole. The bronchial walls are perforated by the openings of numerous alveoli (Fig. 6L). These bronchi and bronchioles have conducting as respiratory function as well.

The lung parenchyma of the adult *Monodelphis domestica* is strongly subdivided and the gas exchange surface area has markedly increased (Tab. 1, Fig. 9D, 10M-P). Short after branching off from the lobar bronchi, the walls of the segmental bronchi are perforated by numerous alveoli (Fig. 6M). These irregular formations of alveoli, similar to that described at 57 dpn, are widespread within the adult lung. An alveolar acinus, typical for the adult mammalian lung, is present. Alveolar sacs contain multiple alveoli, which radiate like a raspberry from the center (Fig. 6N). The alveoli measure ∼50-70 µm in diameter. Compared to earlier developmental stages they have slightly increased in size. In the interalveolar septa pores of Kohn are numerous and present throughout the adult lung. The pores of Kohn (Fig. 6N, arrowheads) form a connection between adjacent alveoli. The single capillary septa separating the alveoli are very thin, they measure 6-10 µm in width. A centrally located capillary occupies the septum almost entirely (Fig. 6O).

Lung volumes (V_L_) and air space volumes (V_A_) in relation to body mass for all specimens examined from neonate to adult are presented in Fig.11A. From neonate to adult a steady increase in lung and air space volume could be observed. Over all, developmental stages, V_L_ (r=0.987) and V_A_ (r=0.915) were closely correlated to body mass.

**Fig. 11.**
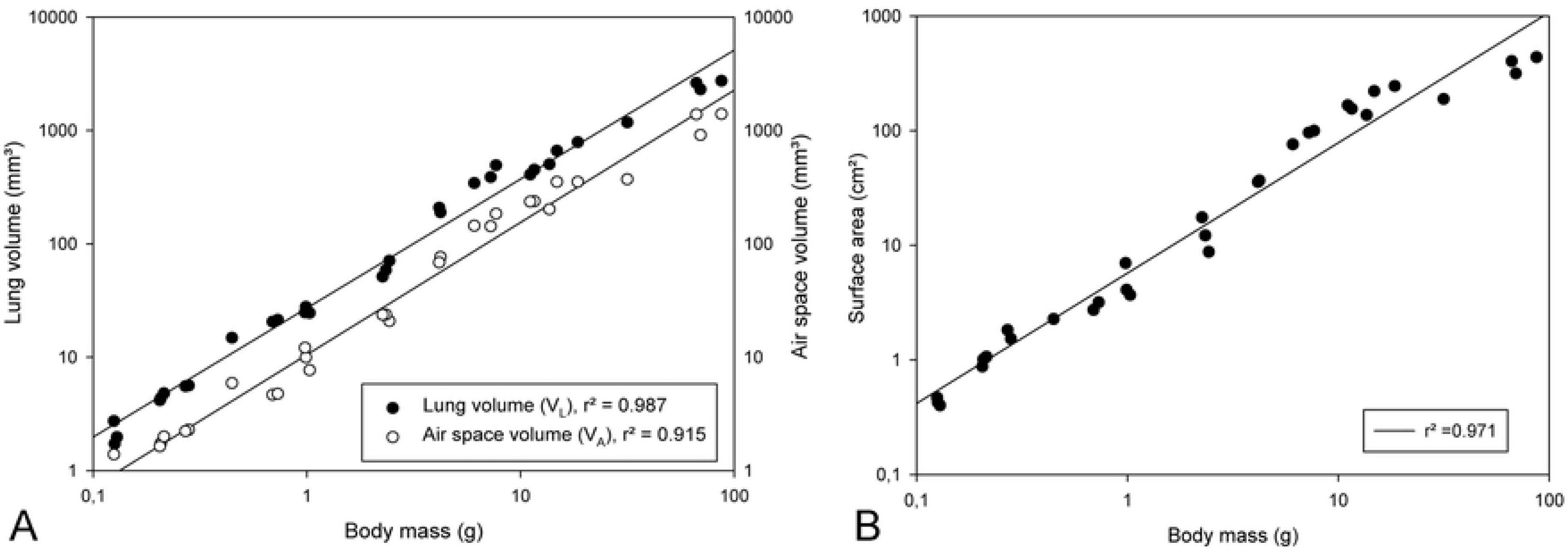
Double logarithmic plots of the lung volume and air space volume (A) and of the surface area (B) against body mass for *Monodelphis domestica* in the postnatal period. The graphs are based on individual animal data and the regression lines are provided.

The surface area (S_A_) of the airspaces in relation to body weight (Fig. 11B) indicates the continuous increase of the gas exchange area during the structural transformation of the lung in the postnatal period. With progressing compartmentalization of the lung by means of subseptation of the lung parenchyma, the airspace surface area increased steadily. The airspace surface area was positively correlated with body mass (r=0.971).

## Discussion

With birth, the lungs of newborn mammals have to take over the function of gas exchange, before provided by the placenta. Viability of the neonate depends on an adequately developed respiratory system (Frappell and Mac Farlane, 2006; Simpson et al., 2013). The lungs of newborn marsupials are not fully developed at birth, as they are born in a relatively immature state compared to placental mammals. Therefore, cutaneous respiration supports gas exchange to various amount depending on the degree of maturity of the lung (Mortola et al., 1999; Ferner, 2018; MacFarlane et al., 2002; Frappell and MacFarlane, 2006).

The lung structure of marsupial neonates follows the size variation in the sequence G1 to G3, proposed by Hughes and Hall (1988). A gradation of lung development from early canalicular stage (G1), late canalicular (G2) to early saccular stage (G3) can be observed among newborn marsupials (Mess and Ferner, 2010; Ferner, 2018). The newborn Gray short-tailed opossum has large terminal airspaces, which are only subdivided a little. The cranial, middle and accessory lobes of the right lung consist of one large terminal airspace respectively. The septum consists of capillaries on both sides, forming a blood air-barrier (Szdzuy et al., 2008) facilitating gas exchange. However, a continuous double capillary septum is not present yet, attributing the lung of the newborn Gray short-tailed opossum to the late canalicular stage (G2). A similar lung structure at birth can be seen in the Virginia opossum (Krause and Leeson, 1975), the brushtail possum (Gemmell and Nelson, 1988) and the bandicoot (Gemmell, 1986; Gemmell and Little, 1982). The most immature lungs among marsupial neonates are present in the dasyurids (G1) like the eastern and northern native cat (Hill and Hill, 1955; Gemmell and Nelson, 1988; Ferner, 2021a) and the stripe-faced and fat-tailed dunnart (Gemmell and Selwood, 1994; Simpson et al., 2011, 2013). They have lungs consisting of well vascularized air chambers that appear like two ‘air bubbles’. More developed lungs (G3) than thus of the newborn Gray short-tailed opossum can be found in kangaroos (Runciman et al., 1996, 1998; Burri et al., 2003; Makanya et al., 2001, 2003, 2007) and the koala (Ferner, 2018). These lungs consist of a primitive bronchial tree terminating in several saccules.

The compartmentalization of the lung of the Gray short-tailed opossum progresses fast during the first postnatal weeks. The terminal airspaces become more and more subdivided by the formation of new double capillary septa from existing septa. With 7 dpn a primary septum with a continuous double capillary bed is present, indicating that the lung has entered the saccular stage. The terminal sacculi become smaller with time and the new formed septa get thinner. The saccular stage can be characterized by the formation of transitory saccules, which are progressively subdivided by septation into more generations of saccules (Ferner et al., 2021a). The process of saccule multiplication is very similar to that of alveolar formation (Burri, 1974; Burri et al., 2003). However, in contrast to the alveolar stage, microvascular maturation, a process that leads to the formation of mature septa with a single capillary layer, does not occur during sacculation.

A prolonged period of saccular subseptation has been described also for the Virginia opossum (Krause and Leeson, 1975), the tammar wallaby (Runciman et al., 1996, 1998), the bandicoot (Gemmel, 1986) and the quokka (Makanya et al., 2001; Burri et al., 2003). Development, from the terminal air sac stage to an alveolar lung, takes place over an extended period of >180 days in the tammar wallaby (Runciman et al., 1998). During this long saccular stage, periods of tissue proliferation and periods of expansion alternate. Up to 20 days, volume increases of the terminal airspaces were largely due to expansion of the air spaces. Around 30 days, rapid tissue proliferation and development of septa which subdivide the air spaces took place. This burst of tissue proliferation was followed by a period during which growth was again largely due to expansion. Between 70 and 180 days subdivision of the large air sacs into much smaller compartments occurred again as the result of tissue proliferation and septal development (Runciman et al., 1996). For the rat a similar differentiation of periods of septa formation and expansion of terminal air spaces has been described during alveolarization (Schittny et al., 2008), indicating analogies in saccular and alveolar formation.

In the Gray short-tailed opossum and most other marsupial species, the saccular stage is much shorter. Alternating periods of tissue proliferation and expansion as described for the tammar wallaby were not detectable. Sacculation in the Gray short-tailed opossum appears to be a continuous process with tissue proliferation and expansion of air spaces taking place in parallel. The first mature single capillary septa, indicating the onset of alveolarization, can be seen at 28 postnatal days and the full alveolar stage is attained at 35 dpn in *Monodelphis domestica*.

Similar times for the onset of alveolarization have been reported for the fat-tailed dunnart (45 dpn; Simpson et al., 2011), the brushtail possum (39 dpn; Buaboocha and Gemmell, 1997), the bandicoot (40 dpn; Gemmell and Little, 1982; Gemmell, 1986) and the Virginia opossum (45 dpn; Krause and Leeson, 1975).

In contrast to marsupials, the lung of placental mammals at birth is either at the late saccular stage in altricial species or at the alveolar stage in precocial species (Szdzuy and Zeller, 2009; Mess and Ferner, 2010). Postnatal lung development in altricial placentals is rapid and the formation of alveoli starts in the golden hamster and mouse at the age of 2 days (Szdzuy et al., 2008; Mund et al., 2008) and in the musk shrew and the rat at 4 days after birth (Burri, 1974; Szdzuy et al., 2008). The majority of placental mammals, among them all precocial species, reach the alveolar stage in utero and possess alveoli already at birth (for review Szdzuy and Zeller, 2009; Mess and Ferner, 2010). The entirety of the numerous small alveoli provides a large gas exchange surface area, a necessity to meet the high metabolic demands accompanying precociality.

The growth of alveoli, including additional formation of new units, proceeds in the postnatal period in marsupial and placental mammals. In marsupials the process of alveolarization continuous until 85 days in the Virginia opossum (Krause and Leeson, 1975), until 113 days in the brushtail possum (Buaboocha and Gemmell, 1997), until 125 days in the quokka (Burri et al., 2003) and until 180 days in the tammar wallaby (Runciman et al., 1996).

For long time it was assumed that alveolarization ceases after the capillary layers in the alveolar septa mature during microvascular maturation (Burri et al., 1974). However, new studies using recently developed techniques report continued alveolarization in placental mammals for rabbits (Kovar et al., 2002), rats (until 60 dpn; Jakkula et al., 2000; Tschanz et al., 2014), and mice (until 36 dpn; Schittny et al., 2008; Mund et al., 2008). Alveolarization continues at least until young adulthood in the rhesus monkey (Hyde et al., 2007) and in humans (Herring et al., 2014; Narayanan et al., 2012). The potential for alveolarization might be preserved throughout life allowing regeneration from degenerative lung diseases (Schittny and Burri, 2007; Schittny, 2017).

New alveolar septa may be lifted off both immature and mature alveolar septa, allowing new septa or new alveoli to be formed principally at any time, even during adulthood (Schittny et al., 2017). Two phases of alveolarization are distinguished: classical (or bulk) alveolarization and continued alveolarization (Schittny et al., 2008; Tschanz et al., 2014). The process of alveolarization has been described in detail by Schittny (2017). The primary septa present at the early saccular stage in marsupials contain a double-layered immature capillary network. At sites where new septa (or secondary septa) will be formed, smooth muscle cell precursors, elastic fibers and collagen fibrils accumulate. The new septa form by an upfolding of one of the two capillary layers. The resulting newly formed secondary septa possess two capillary layers and subdivide preexisting airspaces into smaller sacculi. This process continues as sacculation until the first alveoli are formed. During microvascular maturation, the double-layered capillary network fuses to a single-layered one. During continued alveolarization, new alveolar septa are still formed by an upfolding of the capillary layer, even if the alveolar surface opposing the upfolding is now missing its capillaries. In all modes, sacculation, classical alveolarization and continued alveolarization, a sheetlike capillary layer folds up to form a double-layered capillary network inside the newly formed septum. When a new alveolar septum is formed, it will mature shortly after by a fusion of the double-layered capillary network (Schittny, 2017).

A recent 3D study of the mouse lung suggests that alveolar “septal tips” are in fact ring or purse string structures containing elastin and collagen (Warburton, 2021). Saccular formation and later alveolarization in the terminal airspaces seem to take place by epithelial extrusion through a directionally orientated orifice with a ring or purse string-ring lip that imparts some localized stiffness to the mesenchyme. The alveolar epithelium extrudes outwards into the surrounding mesenchyme, which is correspondingly less stiff than the saccular or alveolar orifice, and forms ‘bubble’ like structures, the sacculi or alveoli (Warburton, 2021).

At birth, the Gray short-tailed opossum weighs approximately 126 mg and has a lung volume (V_L_) of only 1.98 mm³. Adult females have bodyweights of ∼69 g and a V_L_ of 2631.63 mm³. This means that from birth to adulthood the bodyweight increases around 550 times and the V_L_ increase is 1,300 fold. In larger marsupial species an even higher increase of lung volume has been reported, e.g., 3,800 times in the tammar wallaby (Runciman et al., 1998) and 8,000 times in the quokka wallaby (Makanya et al., 2003). In placental mammals the V_L_ increase from birth to adulthood is much lower, only approximately 23 times in humans and rats (Zeltner and Burri, 1987).

Low lung volumes, similar to that of the newborn Gray short-tailed opossum, were reported for other marsupial neonates, e.g., 4 - 9.7 µl in the tammar wallaby (Runciman et al., 1998; Simpson et al., 2013), 2 µl in the quokka (Makanya et al., 2003, 2007) and 0.4 µl in the fat-tailed dunnart (Simpson et al., 2013). For comparison the V_L_ of the newborn rat (body weight 7.2g), a typical altricial placental, is with 570 µl much higher (Makanya et al., 2007). The lung volumes determined in newborn marsupials are lower than that predicted from allometric equations for placental neonates (Frappell and Mac Farlane, 2006). The low lung volume at birth in marsupials may relate to the stage of lung development; the earlier the stage the lower the volume and the greater the extent of cutaneous gas exchange (Simpson et al., 2013).

The volumes of the lung (V_L_) and of the terminal air spaces (V_A_) and the air space surface area (S_A_) increased steadily during the postnatal period in *Monodelphis domestica* (Tab.1). Overall, the developmental stages, V_L_, V_A_ and S_A_ were closely correlated with body mass (Fig. 11). The absolute air space surface area of the newborn lung (0.427 cm²) increased ∼10-fold by day 14 (4.095 cm²). In the more immature born eastern quoll the surface area of the air spaces increased from 0.028 cm² to 2.122 cm² even 26-fold during this time period (Ferner, 2021a). The surface areas of the lung available for gas exchange in the newborn tammar wallaby (1.115 cm²) and dunnart (0.069 cm²), determined by phase contrast X-ray imaging (Simpson et al., 2013), are comparable to the results of this study. Similar to V_L_, the surface areas reported for marsupials are below values predicted from allometry for eutherian species (Frappell and MacFarlane, 2006; Simpson et al., 2013).

Several studies examined morphometric aspects of the developing lung in marsupials (Runciman et al., 1998; Burri et al., 2003; Makanya et al., 2003; Simpson et al., 2013). In the tammar wallaby, studies have shown that changes in the surface area of the lung up to 20 days are largely due to expansion of the air spaces while tissue proliferation and air sac subdivision is most pronounced during the transitional period from ectothermy to endothermy (after 70 postnatal days,100 g) (Runciman et al., 1998). However, in smaller marsupial species, like the Gray short-tailed opossum tissue proliferation and air sac subdivision can be observed already in the early postnatal period and proceeds obviously faster than in the tammar wallaby. A marked increase of air space surface area could be seen in the Gray short-tailed opossum between 14 dpn and 28 dpn, probably due to massive septal development and expansion of existing and new formed sacculi during the late saccular stage. Between 28 and 35 dpn, at the time alveloarization took place, air space surface area more than doubled. In marsupial postnatal development, the airspace surface areas show highest rates of increase during the alveolar stage (Burri et al., 2003; Makanya et al., 2003). However, this correlation of increase in surface area and alveolarization seems to be typical for mammalian lung development in general. Also, morphometric studies in placental species, like the rat, found that the formation of alveoli, occurring between day 4 and 10 results in a five-fold increase in gas exchange surface area (Weibel, 1967; Burri, 1974; Burri et al., 1974). It was found that the alveolar surface area increased by a factor of 2.6 whereas the lung volume increased by only 60 % during this period.

The lung volumes reported in this study might differ from the functional lung volumes of living animals. Limitations due to the method of fixation need to be discussed. Specimens from 0 to 28 dpn were totally immersed in fixative after severing the head to allow fixative inflow. This might result in lower lung volumes than the functional value, because lungs tend to deflate below functional residual capacity (FRC) at death (Simpson et al., 2013). The inflation degree of the lung cannot be controlled with that method, causing a higher variation in lung volumes. In older stages (35dpn - adult) the lungs were filled with fixative to a tracheal pressure of 20 cm. That may lead to an overestimation of lung volume and surface area since lungs inflated with liquid have a larger compliance and are easier to distend than air-filled lungs (West, 1995). Regardless of the preservation methods, the lung volumes and surface areas calculated for *Monodelphis domestica* are comparable to lung and surface areas reported for other marsupial pouch young of similar size. In addition, others have demonstrated the comparability of lung volumes and surface areas derived from computed tomography data sets with histological estimations (Coxon et al., 1999).

## Conclusion

X-ray computed tomography (µCT) offers the possibility to show the comprehensive structural transformations in the developing lung of the Gray short-tailed opossum in 3D. The results confirmed that marsupials such as the Gray short-tailed opossum are born with structurally immature lungs when compared to eutherian mammals and underwent marked increases in architectural complexity during the postnatal period. In marsupials, the process of alveolarization, which takes place largely intrauterine in the eutherian fetus, is shifted to the postnatal period and is therefore more easily accessible for investigation. The structural development of the terminal airspaces from large terminal sacs to the final alveolar lung, takes place in functional state in a continuous morphogenetic process. This allows insights in the structural prerequisites of a functioning lung and opens a new window for better understanding of the evolution of mammalian lung development. It can be assumed that mammalian lung development follows similar developmental pathways in all mammalian species, including marsupials.

## Acknowledgements

I am very grateful to Kristin Mahlow (lab manager of CT-lab) for staining, sample preparation and acquisition of µCT-scans. I thank the animal keepers Petra Grimm and Annett Billepp as well as Dr. Peter Giere and Dr. Peter Bartsch of the animal facility of the Museum für Naturkunde Berlin for breeding and providing the animals for this project.

## Funding

I acknowledge the financial support from the German Research Foundation (DFG) with the module “temporary position for the principal investigator” (Grant No. FE1878/2-1). Open access funding was enabled by the Leibniz Association’s Open Access Publishing Fund and MfN (DFG.Project OA pubication costs).

## Conflict of interest

The authors declare no conflict of interest.

## Data availability statement

The data that support the findings of this study, additional images and videos of 3D reconstructions of the terminal airspaces are available under https://doi.org/10.7479/cy7h-j182.

## Notes

### Competing Interest Statement

The authors have declared no competing interest.

## References

1. Branchfield, K., Li, R., Lungova, V., Verheyden, J. M., McCulley, D., & Sun, X. (2016). A three-dimensional study of alveologenesis in mouse lung. Developmental biology, 409(2), 429–441. 10.1016/j.ydbio.2015.11.017

2. Buaboocha, W., & Gemmell, R. T. (1997). Development of lung, kidney and skin in the brushtail possum, Trichosurus vulpecula. Acta Anatomica, 159, 15–24. 10.1159/000147960

3. Burri, P. H. (1974). The postnatal growth of the rat lung III. Morphology. The Anatomical Record, 180(1), 77–98. 10.1002/ar.1091800109

4. Burri, P. H. (1997). Structural aspects of prental and postnatal development and growth of the lung. In: Growth and Development of the Lung, edited by McDonald J. New York, Basel, Hong Kong: Dekker, p. 1–35.

5. Burri, P. H. (2006). Structural aspects of postnatal lung development–alveolar formation and growth. Neonatology, 89(4), 313–322. 10.1159/000092868

6. Burri, P.H., Dbaly, J., Weibel, E.R. 1974. The postnatal growth of the rat lung. I. Morphometry. The Anatomical Record 178: 711–730. 10.1002/ar.1091780405 PMID: 4592625

7. Burri, P. H., Haenni, B., Tschanz, S. A., & Makanya, A. N. (2003) Morphometry and allometry of the postnatal marsupial lung development: an ultrastructural study. Respiratory physiology & neurobiology, 138(2-3), 309–324. 10.1016/S1569-9048(02)00204-5

8. Coxson, H.O., Rogers, R.M., Whittall, K.P., D’Yachkova, Y., Pare, P.D., et al. (1999) A quantification of the lung surface area in emphysema using computed tomography. American journal of respiratory and critical care medicine 159: 851–856. 10.1164/ajrccm.159.3.9805067

9. Deakin, J. E., Delbridge, M. L., Koina, E., Harley, N., Alsop, A. E., Wang, C.,…&Graves, J. A. M. (2013) Reconstruction of the ancestral marsupial karyotype from comparative gene maps. BMC Evol Biol, 13, 258. 10.1186/1471-2148-13-258

10. Ferner, K. (2018) Skin structure in newborn marsupials with focus on cutaneous gas exchange. Journal of Anatomy, 233(3), 311–327. 10.1111/joa.12843

11. Ferner, K. (2021a) Early postnatal lung development in the eastern quoll (*Dasyurus viverrinus*). The Anatomical Record, 304(12), 2823–2840. 10.1002/ar.24623

12. Ferner, K. (2021b) Development of the skin in the eastern quoll (*Dasyurus viverrinus*) with focus on cutaneous gas exchange in the early postnatal period. Journal of Anatomy, 238(2), 426–445. 10.1111/joa.13316

13. Ferner, K, and Mahlow, K. (2023) 3-D-Reconstruction of the bronchial tree of the Gray short-tailed opossum (*Monodelphis domestica*) in the postnatal period. Journal of Anatomy. Accepted. 10.1111/joa.13928

14. Ferner, K., Zeller, U., & Renfree, M. B. (2009) Lung development of monotremes: evidence for the mammalian morphotype. The Anatomical Record: Advances in Integrative Anatomy and Evolutionary Biology: Advances in Integrative Anatomy and Evolutionary Biology, 292(2), 190–201. 10.1002/ar.20825

15. Ferner, K., Schultz, J.A. & Zeller, U. (2017) Comparative anatomy of neonates of the three major mammalian groups (monotremes, marsupials, placentals) and implications for the ancestral mammalian neonate morphotype. Journal of Anatomy, 231, 798–822. 10.1111/joa.12689

16. Frappell, P. B. & MacFarlane, P. M. (2006) Development of the respiratory system in marsupials. Respiratory Physiology and Neurobiology, 154, 252–267. 10.1016/j.resp.2006.05.001

17. Frappell, P.B. & Mortola, J.P. (2000) Respiratory function in a newborn marsupial with skin gas exchange. Respiration Physiology, 120(1), 35–45. 10.1016/S0034-5687(99)00103-6

18. Gemmell, R.T. (1986) Lung development in the marsupial bandicoot, *Isoodon macrourus*. Journal of Anatomy, 148, 193–204. www.ncbi.nlm.nih.gov/pmc/articles/PMC1261602/

19. Gemmell, R.T. & Little, G.J. (1982) The structure of the lung of the newborn marsupial bandicoot, *Isoodon macrourus*. Cell and Tissue Research, 223, 445–453. 10.1007/BF01258501

20. Gemmell, R.T. & Nelson, J. (1988) The ultrastructure of the lung of two newborn marsupial species, the northern native cat, *Dasyurus hallucatus*, and the brushtail possum, Trichosurus vulpecula. Cell Tissue Research, 252, 683–685. 10.1007/BF00216657

21. Gemmell RT, Selwood L (1994) Structural development in the newborn marsupial, the Stripe-faced dunnart, Sminthopsis macroura. Acta Anat 149, 1–12. 10.1159/000147549

22. Gignac, P. M., Kley, N. J., Clarke, J. A., Colbert, M. W., Morhardt, A. C., Cerio, et al. (2016). Diffusible iodine-based contrast-enhanced computed tomography (diceCT): an emerging tool for rapid, high-resolution, 3-D imaging of metazoan soft tissues. Journal of Anatomy, 228(6), 889-909. 10.1111/joa.12449. Epub 2016 Mar 11. PMID: 26970556; PMCID: PMC5341577.

23. Haberthür, D., Yao, E., Barré, S.F., Cremona, T.P., Tschanz, S.A., Schittny, J.C. (2021). Pulmonary acini exhibit complex changes during postnatal rat lung development. PLoS ONE 16(11): e0257349. 10.1371/journal.pone.0257349

24. Herring, M.J., Putney, L.F., Wyatt, G., Finkbeiner, W.E., Hyde, D.M. (2014) Growth of alveoli during postnatal development in humans based on stereological estimation. Am J Physiol Lung Cell Mol Physiol 307:L338–L344. 10.1152/ajplung.00094.2014

25. Hill, J.P. & Hill, W.C.O. 1955 The growth stages of the pouch young of the native cat (Dasyurus viverrinus) together with observations on the anatomy of the newborn young. Transactions of the Zoological Society of London, 28, 349–352. 10.1111/j.1096-3642.1955.tb00003.x

26. Hughes, R.L., Hall, L.S. (1988). Structural Adaptations of the Newborn Marsupial. In: Tyndale-Biscoe, C.H., Janssens, P.A. (eds) The Developing Marsupial. pp. 8–27. Springer, Berlin, Heidelberg. 10.1007/978-3-642-88402-3_2

27. Hyde, D.M., Blozis, S.A., Avdalovic, M.V., Putney, L.F., Dettorre, R., Quesenberry, N.J., Singh, P., Tyler, N.K. (2007). Alveoli increase in number but not size from birth to adulthood in rhesus monkeys. Am J Physiol Lung Cell Mol Physiol 293: L570–L579. 10.1152/ajplung.00467.2006

28. Jakkula, M., Le Cras, T.D., Gebb, S., Hirth, K.P., Tuder, R.M., Voelkel, N.F., Abman, S.H. (2000). Inhibition of angiogenesis decreases alveolarization in the developing rat lung. Am J Physiol Lung Cell Mol Physiol 279:L600–L607. 10.1152/ajplung.2000.279.3.L600

29. Jeffery, P.K. (1998). The development of large and small airways. American Journal of Respiratory and Critical care Medicine, 157(5), S174–S180. 10.1164/ajrccm.157.5.rsaa-1

30. Kitaoka, H., Burri, P. H., & Weibel, E. R. (1996). Development of the human fetal airway tree: analysis of the numerical density of airway endtips. The Anatomical Record: An Official Publication of the American Association of Anatomists, 244(2), 207–213. 10.1002/(SICI)1097-0185(199602)244:2<207::AID-AR8>3.0.CO;2-Y

31. Kovar, J., Sly, P.D., Willet, K.E. (2002). Postnatal alveolar development of the rabbit. J Appl Physiol 93:629–635. 10.1152/japplphysiol.01044.2001

32. Krause, W.J. & Leeson, C.R. (1975). Postnatal development of the respiratory system of the opossum. II. Electron microscopy of the epithelium and pleura. Acta Anatomica, 92, 28–44. 10.1159/000144426

33. MacFarlane, P.M. & Frappell, P.B. (2001) Convection requirement is established by total metabolic rate in the newborn tammar wallaby. Respiration Physiology, 126, 221–231. 10.1016/S0034-5687(01)00227-4

34. MacFarlane, P. M., Frappell, P. B., & Mortola, J. P. (2002). Mechanics of the respiratory system in the newborn tammar wallaby. Journal of Experimental Biology, 205(4), 533–538. 10.1242/jeb.205.4.533

35. Makanya, A.N., Sparrow, P., Warui, N., Mwangi, K. & Burri, H. (2001) Morphological analysis of the postnatally developing marsupial lung: the quokka wallaby. The Anatomical Record: An Official Publication of the American Association of Anatomists, 262(3), 253–265. 10.1002/1097-0185(20010301)262:3<253:AID-AR1025>3.0.CO;2-B

36. Makanya, A.N., Haenni, B. & Burri, P.H. (2003) Morphometry and allometry of the postnatal lung development in the quokka wallaby (*Setonix brachyurus*): a light microscopic study. Respiratory Physiology & Neurobiology, 134(1), 43–55. 10.1016/S1569-9048(02)00204-5

37. Makanya, A.N., Tschanz, S.A., Haenni, B. & Burri, P.H. (2007) Functional respiratory morphology in the newborn quokka wallaby (*Setonix brachyurus*). Journal of Anatomy, 211, 26–36. 10.1111/j.1469-7580.2007.00744.x

38. Mess, A.M. & Ferner, K.J. (2010) Evolution and development of gas exchange structures in Mammalia: the placenta and the lung. Respiratory Physiology & Neurobiology,173, 74–82. 10.1016/j.resp.2010.01.005

39. Metscher, B.D. (2009) MicroCT for comparative morphology: simple staining methods allow high-contrast 3D imaging of diverse non-mineralized animal tissues. BMC Physiology, 9:11. 10.1186/1472-6793-9-11. PMID: 19545439; PMCID: PMC2717911.

40. Miller, N. J., Orgeig, S., Daniels, C. B., & Baudinette, R. V. (2001). Postnatal development and control of the pulmonary surfactant system in the tammar wallaby Macropus eugenii. Journal of Experimental Biology, 204(23), 4031–4042. 10.1242/jeb.204.23.4031

41. Modepalli, V., Kumar, A., Sharp, J. A., Saunders, N. R., Nicholas, K. R. & Lefèvre, C. (2018) Gene expression profiling of postnatal lung development in the marsupial gray short-tailed opossum (*Monodelphis domestica*) highlights conserved developmental pathways and specific characteristics during lung organogenesis. BMC genomics, 19(1), 732. 10.1186/s12864-018-5102-2

42. Morrisey, E. E., & Hogan, B. L. (2010). Preparing for the first breath: genetic and cellular mechanisms in lung development. Developmental cell, 18(1), 8–23. 10.1016/j.devcel.2009.12.010

43. Mortola, J. P., Frappell, P. B., & Woolley, P. A. (1999). Breathing through skin in a newborn mammal. Nature, 397(6721), 660. 10.1038/17713

44. Mulisch, M., & Welsch, U. (Eds.). (2015) Romeis-Mikroskopische Technik. Springer-Verlag. Berlin, Heidelberg.

45. Mund, S. I., Stampanoni, M., & Schittny, J. C. (2008). Developmental alveolarization of the mouse lung. Developmental dynamics: an official publication of the American Association of Anatomists, 237(8), 2108–2116. 10.1002/dvdy.21633

46. Narayanan, M., Owers-Bradley, J., Beardsmore, C.S., Mada, M., Ball, I., Garipov, R., Panesar, K.S., Kuehni, C.E., Spycher, B.D., Williams, S.E., Silverman, M. (2012). Alveolarization continues during childhood and adolescence: new evidence from helium-3 magnetic resonance. Am J Respir Crit Care Med 185:186–191. 10.1164/rccm.201107-1348OC

47. Post, M., & Copland, I. (2002). Overview of lung development. Acta Pharmacologica Sinica, 23(SUPP), 4–7. 10.1016/S1526-0542(04)90049-8

48. Ribbons, K. A., Baudinette, R. V., & McMurchie, E. J. (1989). The development of pulmonary surfactant lipids in a neonatal marsupial and the rat. Respiration physiology, 75(1), 1–10. 10.1016/0034-5687(89)90081-9

49. Renfree, M.B. (2006) Society for Reproductive Biology Founders’ Lecture 2006. Life in the pouch: womb with a view. Reproduction Fertility and Development 18(7), 721–734. 10.1071/RD06072

50. Runciman, S.I.C., Gannon, B.J. & Baudinette, R.V. (1995). Central cardiovascular shunts in the perinatal marsupial. Anatomical Record, 243(1), 71–83. 10.1002/ar.1092430109

51. Runciman, S.I.C., Baudinette, R.V. & Gannon, B.J. (1996). Postnatal development of the lung parenchyma in a marsupial: the tammar wallaby. Anatomical Record, 244, 193–206. 10.1002/(SICI)1097-0185(199602)244:2<193::AID-AR7>3.0.CO;2-2

52. Runciman, S.I.C., Baudinette, R.V., Gannon, B.J. & Lipsett, J. (1998) Morphometric analysis of postnatal lung development in the tammar wallaby: light microscopy. Respiration Physiology, 112, 325–337. 10.1016/S0034-5687(98)00034-6

53. Schittny, J.C. (2017) Development of the lung. Cell and Tissue Research, 367, 427–444. 10.1007/s00441-016-2545-0

54. Schittny JC, Burri PH. 2007. Development and growth of the lung. In: Fishman AP, Elias JA, Fishman JA, Grippi MA, Kaiser R, Senior RM, eitors. Fishman’s pulmonary diseases and disorders. New-York: McGraw-Hill.

55. Schittny, J. C., Mund, S. I., & Stampanoni, M. (2008). Evidence and structural mechanism for late lung alveolarization. American Journal of Physiology-Lung Cellular and Molecular Physiology, 294(2), L246–L254. 10.1152/ajplung.00296.2007

56. Simpson, S.J., Flecknoe, S.J., Clugston, R.D., et al. (2011) Structural and functional development of the respiratory system in a newborn marsupial with cutaneous gas exchange. Physiological and Biochemical Zoology, 84(6), 634–649. 10.1086/662557

57. Simpson, S.J., Siu, K.K., Yagi, N., Whitley, J.C., Lewis, R.A. & Frappell, P.B. (2013) Phase contrast imaging reveals low lung volumes and surface areas in the developing marsupial. Plos one, 8(1), e53805. 10.1371/journal.pone.0053805

58. Szdzuy, K. (2006). Reproductive strategies of K/t-crossing Theria - Neonate and postnatal development of the morphotype of Marsupialia and Placentalia (Mammalia). Diss. Humbodt-Universität. Berlin. 10.18452/15483

59. Szdzuy, K. & Zeller, U. (2009). Lung and metabolic development in mammals: contribution to the reconstruction of the marsupial and eutherian morphotype. Journal of Experimental Zoology, 312B, 555–578. 10.1002/jez.b.21228

60. Szdzuy, K., Zeller, U., Renfree, M., Tzschentke, B. & Janke, O. (2008) Postnatal lung and metabolic development in two marsupial and four eutherian species. Journal of Anatomy, 212(2), 164–179. 10.1111/j.1469-7580.2007.00849.x

61. Ten Have-Opbroek, A. A. (1981). The development of the lung in mammals: an analysis of concepts and findings. American Journal of Anatomy, 162(3), 201–219. 10.1002/aja.1001620303

62. Tschanz, S.A. (2007). Structural aspects of pre-and post-natal lung development. Pneumologie, 61(7), 479–481. 10.1055/s-2007-959221

63. Tschanz, S.A., Salm, L.A., Roth-Kleiner, M., Barré, S.F., Burri, P.H., Schittny, J.C. (2014). Rat lungs show a biphasic formation of new alveoli during postnatal development. Journal of Applied Physiology 117: 89–95. 10.1152/japplphysiol.01355.2013 PMID: 24764134

64. Vasilescu, D.M., Knudsen, L., Ochs, M., Weibel, E.R. & Hoffman, E.A. (2012). Optimized murine lung preparation for detailed structural evaluation via micro-computed tomography. Journal of Applied Physiology, 112(1), 159–166. 10.1152/japplphysiol.00550.2011

65. Wang, Z., Hubbard, G.B., Clubb Jr, F.J. & VandeBerg, J.L. (2009). The laboratory opossum (*Monodelphis domestica*) as a natural mammalian model for human cancer research. International Journal of Clinical and Experimental Pathology, 2(3), 286. PMID: 19079623; PMCID: PMC2600460

66. Warburton, D. (2021). Conserved Mechanisms in the Formation of the Airways and Alveoli of the Lung. Frontiers in Cell and Developmental Biology, 9, 662059. 10.3389/fcell.2021.662059

67. Warburton, D., El-Hashash, A., Carraro, G., Tiozzo, C., Sala, F., et al. (2010). Lung organogenesis. Current Topics in Developmental Biology, 90, 73–158. 10.1016/S0070-2153(10)90003-3.

68. Weibel, E. R. (1967). Postnatal growth of the lung and pulmonary gas-exchange capacity. In R. Porter (Ed.), de Reuck AVS (pp. 131–148). Development of the Lung. A CIBA Foundation Symposium. London: Churchill. 10.1002/9780470719473.ch8

69. West, J.B. (1995). Mechanics of breathing. Respiratory physiology – the essentials. Pp. 89–116. Williams & Wilkins. Baltimore

70. Zeltner, T.B. & Burri, P.H. (1987). The postnatal development and growth of the human lung. II. Morphology. Respir Physiol 67, 269–282. 10.1016/0034-5687(87)90058-2

